# A *Vibrio cholerae* Anti-Phage System Depletes Nicotinamide Adenine Dinucleotide to Restrict Virulent Bacteriophages

**DOI:** 10.1101/2024.06.17.599363

**Authors:** Yishak A. Woldetsadik, David W. Lazinski, Andrew Camilli

**Affiliations:** Department of Molecular Biology and Microbiology, Graduate School of Biomedical Sciences, Tufts University School of Medicine, Boston, Massachusetts, USA

## Abstract

Bacteria and their predatory viruses (bacteriophages or phages) are in a perpetual molecular arms race. This has led to the evolution of numerous phage defensive systems in bacteria that are still being discovered, as well as numerous ways of interference or circumvention on the part of phages. Here, we identify a unique molecular battle between the classical biotype of *Vibrio cholerae* and virulent phages ICP1, ICP2, and ICP3. We show that classical biotype strains resist almost all isolates of these phages due to a 25-kb genomic island harboring several putative anti-phage systems. We observed that one of these systems, Nezha, encoding SIR2*-*like and helicase proteins, inhibited the replication of all three phages. Bacterial SIR2-like enzymes degrade the essential metabolic coenzyme nicotinamide adenine dinucleotide (NAD^+^), thereby preventing replication of the invading phage. In support of this mechanism, we identified one phage isolate, ICP1_2001, which circumvents Nezha by encoding two putative NAD^+^ regeneration enzymes. By restoring the NAD^+^ pool, we hypothesize that this system antagonizes Nezha without directly interacting with either protein and should be able to antagonize other anti-phage systems that deplete NAD^+^.

## Introduction

*Vibrio cholerae* is a gram-negative facultative pathogen that resides in estuarine environments and causes the severe diarrheal disease cholera. There are two biotypes of *V. cholerae*: classical, which caused the first six recorded cholera pandemics, and El Tor, responsible for the ongoing 7^th^ pandemic (1). In areas surrounding the Bay of Bengal, *V. cholerae* is frequently preyed upon within the intestinal tract of cholera victims by three virulent phages: ICP1, ICP2, and ICP3. ICP1 is the most commonly isolated of the three phages both in cholera patient stools and the aquatic environment (2, 3) and is the only one of the three that can prey on *V. cholerae* in the aquatic environment (4). Thus, *V. cholerae* and ICP1 frequently engage in a co-evolutionary arms race. As a result, *V. cholerae* possesses several lines of defense against ICP1, some appearing universal in the species and some varying from strain to strain. One universal mechanism is the phase variation of the lipopolysaccharide O1-antigen, which is the receptor for ICP1 (5). Other defense systems are cytoplasmically located and include diverse mechanisms that interfere with various stages of the ICP1 life cycle, such as phage-inducible chromosomal island-like elements (PLEs) present in many classical and El Tor strains and the cyclic-oligonucleotide-based anti-phage signaling system (CBASS) found in all El Tor strains (6, 7).

The PLE excises from the chromosome upon ICP1 infection and blocks phage replication by degrading the phage DNA and causing abortive infection by premature lysis. PLEs can spread horizontally by hijacking ICP1 structural components to make PLE- transducing virions (8). Some ICP1 isolates overcome this potent defense system by encoding a Type I-F CRISPR/Cas system or a standalone nuclease, both targeting the PLE (9). Classical biotype strains appear to have only one version of the PLE, PLE5, whereas El Tor strains may have one of several different PLE types, though not PLE5 (10). Additionally, the El Tor biotype lacks a CRISPR/Cas system, whereas classical strains encode a Type 1-E CRISPR/Cas system for phage defense (11, 12). Given the differences in phage defensive systems between classical and El Tor biotypes, we asked if there are biotype differences in sensitivity to ICP1 infection. Here, we use a collection of classical biotype strains and ICP1 isolates to show that classical biotype strains are almost universally resistant to ICP1 independent of CRISPR/Cas. Based on this, we hypothesized that classical biotype strains encode an unknown anti-phage system that restricts ICP1.

To uncover this anti-phage system, we used whole genome sequencing and bioinformatics to identify two putative anti-phage systems, Gabija and Nezha, residing on a 25-kb genetic element unique to the classical biotype. These two systems were recently characterized in other bacteria and use distinct mechanisms for subverting phages. Gabija encodes for a protein complex composed of a GajA nuclease and GajB helicase (13, 14). Gabija is activated by NTP depletion, resulting in the selective degradation of DNA in phage-infected cells (15). Nezha encodes for a protein complex composed of a SIR2-like Nicotinamide adenine dinucleotide (NAD^+^) hydrolase and a HerA helicase (16–18). Nezha depletes the essential metabolic coenzyme NAD^+^, thus blocking phage replication (16, 17, 19). Here, we show Nezha is responsible for conferring resistance to ICP1, ICP2, and ICP3. Highlighting the consequences of phage-host coevolution, we identify one ICP1 isolate that counters Nezha using a NAD^+^ regenerating system. Our results provide insight into a previously uncharacterized host-phage arms race in classical biotype *V. cholerae*.

## Results

### Classical biotype *V. cholerae* encodes anti-phage systems absent in the El Tor biotype

Using a collection of 12 classical biotype *V. cholerae* clinical strains isolated from multiple countries between 1949 and 1970, we found that all were resistant to infection by the modern virulent *Myoviridae* phage ICP1_2011_A. In contrast, eight El Tor biotype clinical isolates from 1978-2022 were sensitive to this phage. Although we lack ICP1 isolates from 1949-1970, we infer this phage was present due to the ICP1-specific PLE5 immunity system in classical biotype strains (10). ICP1_2011_A blocks PLE5 immunity by encoding a CRISPR/Cas system with a spacer targeting PLE5 (9). To further probe this biotype difference in phage resistance, we tested representative classical and El Tor biotype strains A103 and E7946, respectively, for sensitivity to a panel of ICP1 isolates from 2001-2011. A103 was resistant to all ICP1 phages except one, ICP1_2001, whereas E7946 was sensitive to all **(Table 1)**. Based on this, we hypothesize that classical biotype *V. cholerae* possesses a phage defensive system absent from El Tor and that ICP1_2001 contains gene(s) absent from the other ICP1 isolates that allow it to overcome the phage defensive system.

**Table 1.** Phage sensitivity profile of representative classical and El Tor biotype strains against ICP1 phage isolates. (+) indicates sensitivity, while (-) indicates resistance to ICP1.

|  |  | Phage isolates |  |  |  |  |  |  |  |
| --- | --- | --- | --- | --- | --- | --- | --- | --- | --- |
|  |  | ICP1_2001 | ICP1_2001_A | ICP1_2004_A | ICP1_2005_A | ICP1_2006_C | ICP1_2006_D | ICP1_2006_E | ICP1_2011_A |
| Strain biotype | E7946 El Tor | + | + | + | + | + | + | + | + |
|  | A103 classical | + | - | - | - | - | - | - | - |

We used whole-genome sequencing and bioinformatic analysis to identify differences between the two *V. cholerae* biotypes concerning phage resistance. We scanned the genomes for anti-phage systems using the bioinformatics program DefenseFinder (20–22). We found a 25-kb genomic island in all 12 classical biotype strains that was absent in El Tor, located between the common genes *tonB* and *trmA* **(Figure 1)**. This island encodes several putative anti-phage systems, including Gabija (*gajAB*) and Nezha (*SIR2-herA*). Based on this, we hypothesized that Gabija and/or Nezha contribute to the phage resistance phenotype observed for classical biotype strains.

**Figure 1:**
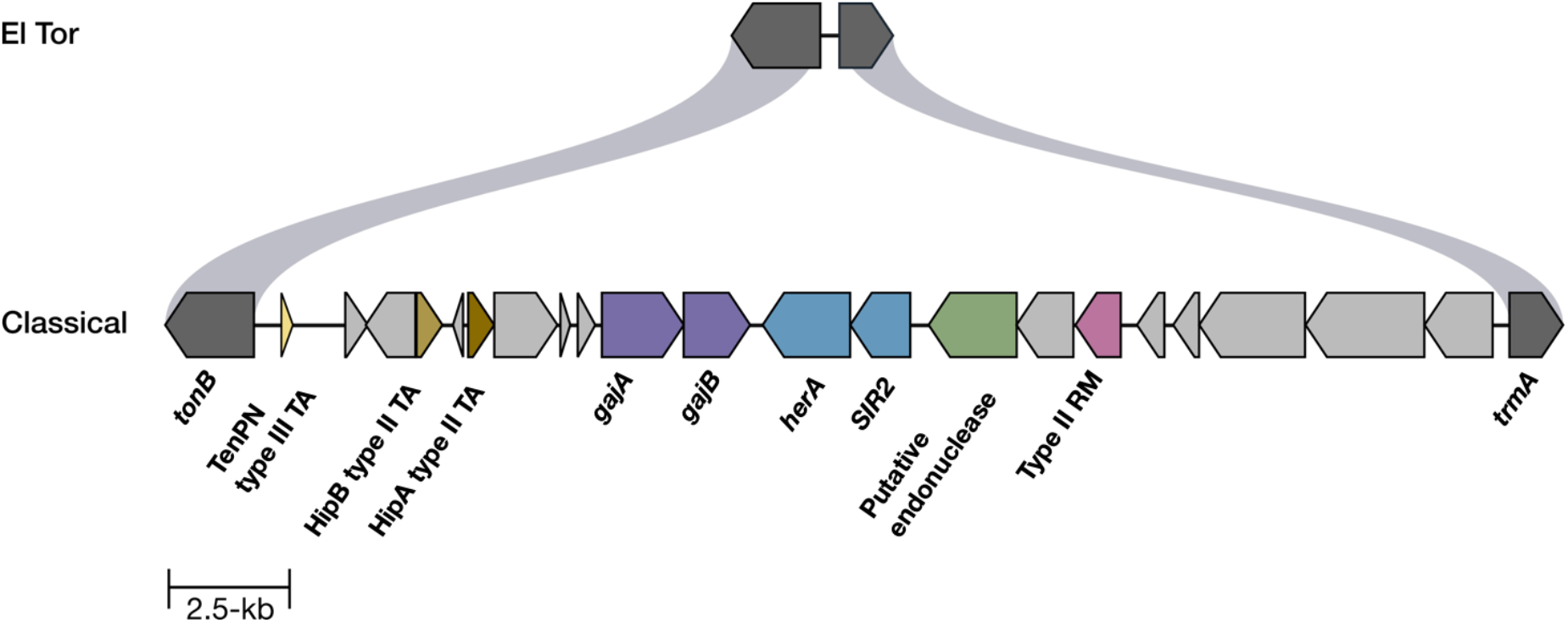
Classical biotype *V. cholerae* encode a genomic island with phage immunity genes. Schematic showing the genome organization of a unique 25-kb genomic island flanked by tonB and trmA in classical *V. cholerae* strain A103. From left to right, the island contains several putative anti-phage systems, including a type III toxin-antitoxin system, type II toxin-antitoxin system, Gabija (*gajAB*), Nezha (*SIR2-herA*), an endonuclease, and a type II restriction-modification system. Genes in gray are of unknown function.

### Nezha provides immunity against ICP1

We generated marked gene deletions using an apramycin resistance cassette to test the role of the whole island, Gabija, and Nezha in the classical biotype strain A103. Five deletion strains were made: Δisland, a double system deletion *ΔΔ* (*ΔgajA ΔgajB* and *ΔSIR2 ΔherA*), the single system deletions (*ΔgajA ΔgajB* or *ΔSIR2 ΔherA*), and deletion of all genes in the island except for *SIR2 herA* **(Figure 2A)**. These and other mutants constructed in this study were whole genome sequenced to confirm the desired mutations and lack of other mutations. The role of Gabija or Nezha in phage resistance was tested by plaque assays on the wild type or mutant A103 derivatives using soft agar overlays containing individual *V. cholerae* strains and spotting ten-fold serial dilutions of the phages indicated. As expected, the wild type A103 strain was sensitive to ICP1_2001 but resistant to ICP1_2001_A and ICP1_2004_A **(Figure 2A, B).** However, the Δisland and the *ΔΔ* mutants were sensitive to all three phages, suggesting that Gabija and/or Nezha mediate phage resistance. Assaying the single system deletion strains revealed that Nezha (*SIR2 herA*) is the primary anti-phage system and that Gabija plays only a minor role in resistance. This was confirmed by showing that deletion of the entire island except for *SIR2 herA* retained resistance to ICP1_2001_A and ICP1_2004_A **(Figure 2A, B)**. Overall, these results show that most ICP1 isolates are blocked by Nezha.

**Figure 2.**
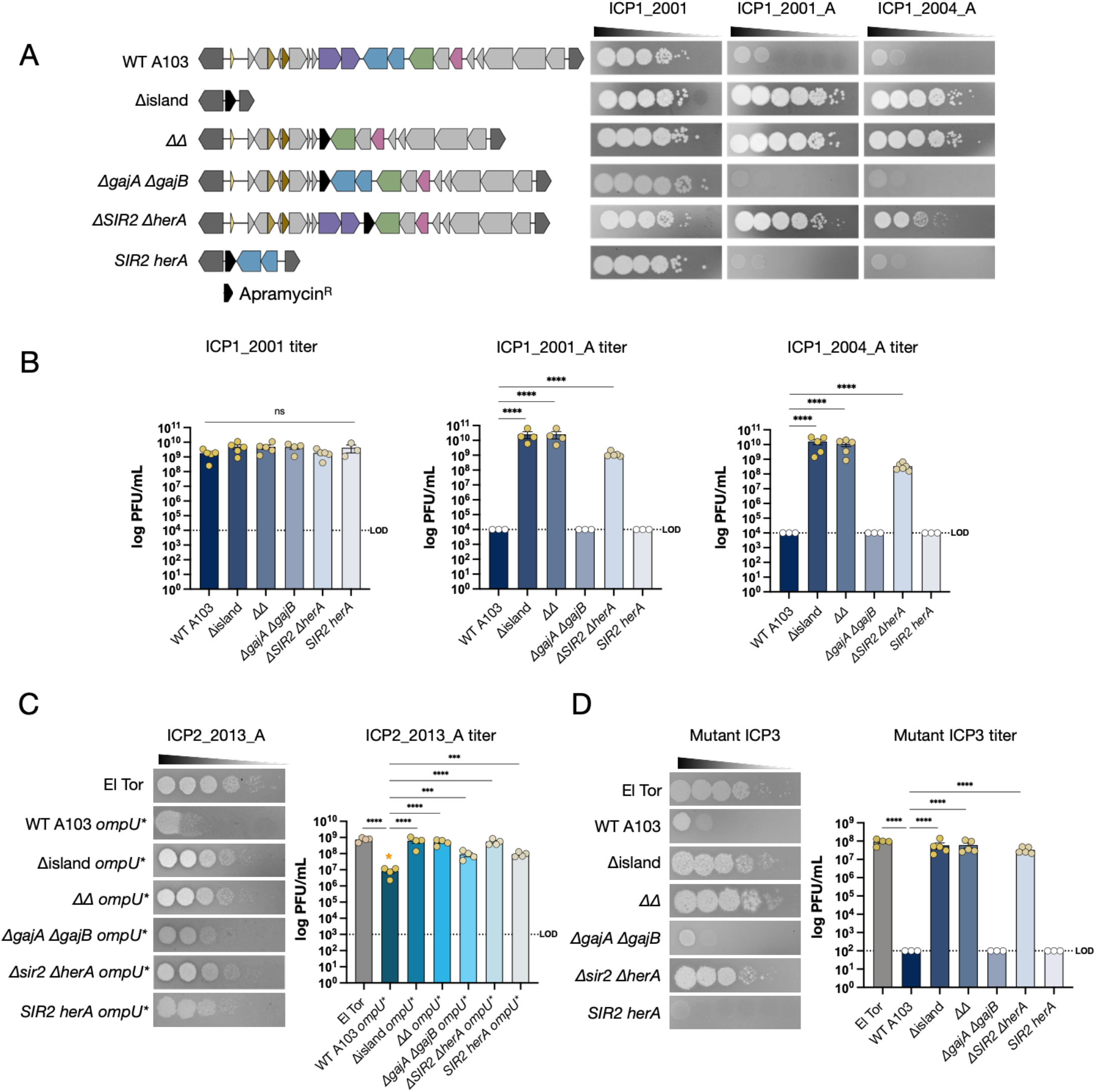
Sir2-HerA protects classical biotype *V. cholerae* from several phages. **(A)** Schematic of the gene deletions constructed in the classical biotype strain A103 (left) and ICP1 phage sensitivity profiles (right). The phage sensitivity of each strain on the left is shown by plaque formation after spotting ten-fold serial dilutions of the ICP1 phages indicated at the top. ICP1_2001 and ICP1_2001_A are distinct isolates from the same year. The results from 3-5 biological replicate experiments are quantified in **(B)**. White-filled circles indicate titers below the limit of detection (LOD) indicated by the horizontal dashed lines. **(C)** Phage sensitivity of the strains indicated to phage ICP2_2013_A (left) and quantification of the results from 4 biological replicates (right). El Tor is strain E7946; the rest are classical strain A103 wild type and derivatives. Because the gene for the ICP2 receptor, *ompU*, is naturally mutated in A103, it was replaced with the El Tor E7946 *ompU* (indicated by *ompU*\*). The orange asterisk (*) indicates incubation for 3 hrs at 37°C. **(D)** Phage sensitivity of the strains indicated to mutant ICP3 phage (left) and quantification of the results from 3-5 biological replicates (right). Data are shown as the standard error of the mean. Analysis was performed using one-way ANOVA with Dunnett’s multiple comparison test (*\*P<0.05, **P<0.01, ***P<0.001, ****P<0.0001*).

### Nezha provides immunity against ICP2 and ICP3

We tested whether Nezha protects against other phages. In contrast to ICP1 and the *Podoviridae* phage ICP3, which both use the lipopolysaccharide O1-antigen as their receptors (2, 23), the distinct *Podoviridae* phage ICP2 uses the outer membrane porin OmpU as its receptor (24). Work done using the El Tor biotype showed that amino acid changes at positions V324 and G325 within an outer loop of OmpU confer resistance to ICP2 (24, 25). In the classical strain A103, OmpU naturally varies at both positions (V324A and G325S), leading to ICP2 resistance in both the wild type and Δisland strains **(Supplementary Figure 1)**. To overcome this block, we transformed naturally competent A103 and its derivatives to replace *ompU* with that from El Tor strain E7946 (designated *ompU\** in Figure 2C). Classical strain A103 with El Tor *ompU* (WT OmpU*) was partially sensitive to ICP2, as small plaques formed after 3 hours at 37°C at the two highest phage concentrations but then disappeared after 4 hours. In contrast, infecting both Δisland OmpU* and *ΔΔ* OmpU* strains yielded complete sensitivity to ICP2 with plaquing efficiencies comparable to that on the El Tor E7946 positive control strain **(Figure 2C)**. Once again, Nezha appears to be the primary resistance determinant, as shown by a significant 100-fold increase in plaquing efficiency in *ΔSIR2 ΔherA* OmpU* compared to WT OmpU*, which was comparable to the plaquing efficiencies on the Δisland OmpU* and *ΔΔ* OmpU* strains.

Classical strain A103 was also resistant to infection by ICP3 **(Figure 2D)**. We do not believe this resistance is due to receptor availability since the classical and El Tor biotypes express the same O1-antigen. In contrast to ICP1 and ICP2, deleting Nezha and Gabija or the entire island did not allow for plaque formation by ICP3. We hypothesized that classical strain A103 has an additional phage defensive system against ICP3 that obscures Nezha and Gabija’s potential role. To circumvent this putative additional ICP3 defense system, we sought to isolate spontaneous mutants of ICP3 that could form plaques on the Δisland strain. We readily obtained ICP3 mutants that formed plaques on the A103 Δisland strain. Sequencing two independent pairs of mutants revealed that each pair had a frameshift mutation in different locations within *orf17* **(Supplemental Table 1).** Orf17 lacks detectable sequence or predicted structural homology to proteins of known function, so the nature of the host immune mechanism circumvented by these null mutations is unknown. Regardless, the mutant ICP3 allowed us to probe the roles of Nezha and Gabija. Upon infecting a *ΔgajA ΔgajB orf17* strain, we observed no plaques, which suggests that Gabija does not inhibit ICP3. However, infection of a *ΔSIR2 ΔherA orf17* strain rescued plaque formation to a level equivalent to that on the Δisland and El Tor E7946 strains (**Figure 2D**). This indicates that Nezha, but not Gabija, antagonizes ICP3. These data thus far show that three different phages, ICP1, ICP2, and ICP3, are antagonized by the Nezha (*SIR2 herA*) anti-phage system, with Gabija (*gajAB*) playing a minor role in resistance to ICP1_2001_A, ICP1_2004_A, and ICP2.

### Mutating the active site of SIR2 or deleting *herA* abolishes phage immunity

The Nezha homologs SIR2 and HerA from *Escherichia coli* form a large protein complex that provides phage immunity through multiple mechanisms, including NAD^+^ hydrolysis, helicase, and nuclease activities (16, 17). However, other studies on homologous systems showed that point mutations in the conserved NADase active site of SIR2 is sufficient to abolish immunity, suggesting that depletion of NAD^+^ is critical for Nezha- mediated phage defense (16, 17, 19). Amino acid sequence and protein structure alignments to *E. coli* SIR2 indicate that N168 and H227 of A103 SIR2 are conserved NADase active site residues **(Figure 3A)**. To examine whether the NADase activity of the A103 SIR2 protein is important for phage defense, we generated alanine substitutions in both active site residues in the *ΔgajA ΔgajB* mutant strain background **(Figure 3A)**. Active site mutations N168A or H227A abolished immunity to ICP1, ICP2, and ICP3 **(Figure 3B-F)**. Since HerA forms a complex with SIR2, it may be needed for SIR2 function. To test this, we deleted *herA* and measured phage sensitivity. We observed that ICP1_2001_A, ICP1_2004_A, and ICP3 had significantly higher plaquing efficiencies on the *ΔherA* strain when compared to WT A103, indicating that HerA complex formation with SIR2 is required for phage immunity **(Supplemental Figure 2)**.

**Figure 3.**
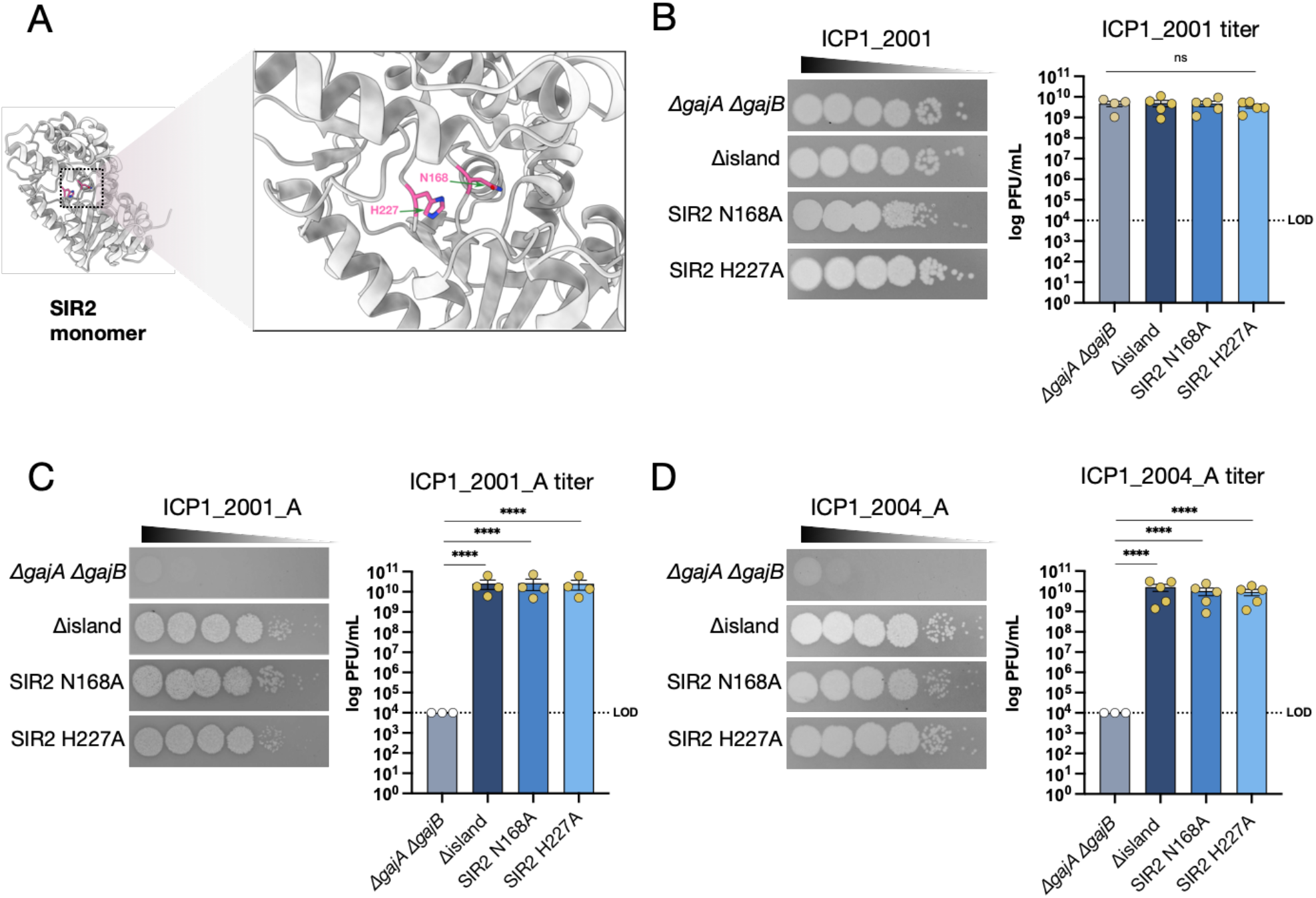
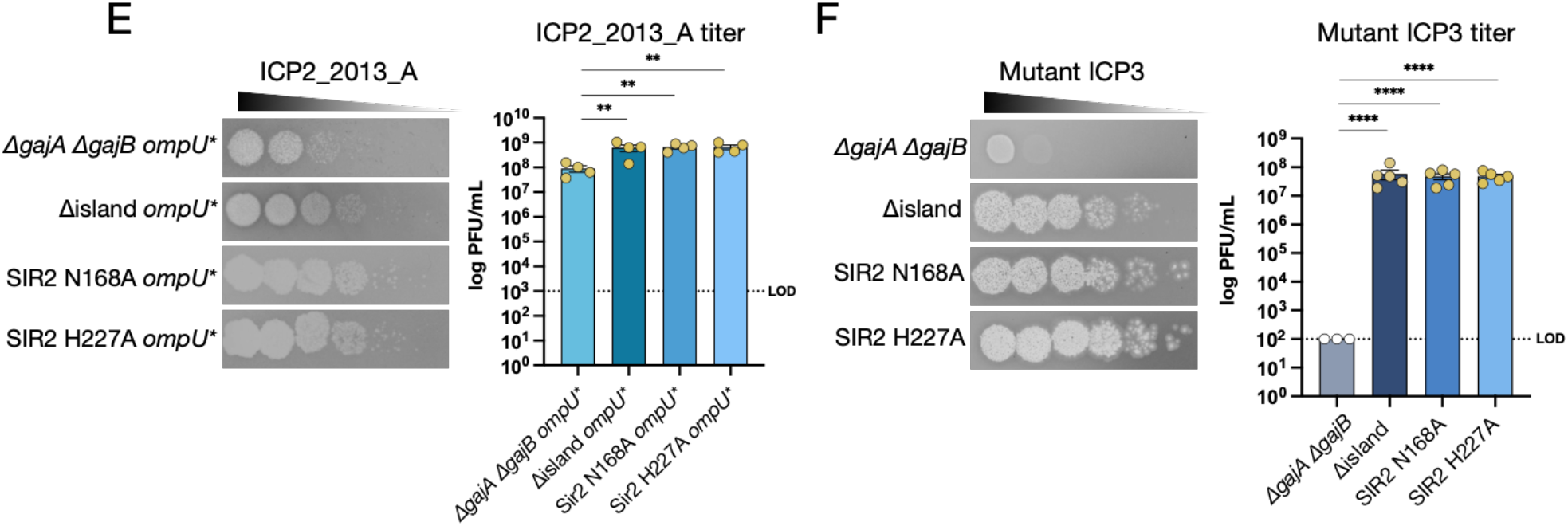
Mutation of the SIR2 NADase active site abolishes immunity to ICP1 and ICP3. **(A)** The SIR2 protein monomer’s structure predicted by Alphafold (18) showing active site residues N168A and H227A (24). **(B-D)** Phage sensitivity of the strains indicated to ICP1 phages (left) and quantification of the results from 3-5 biological replicates (right). **(E)** Phage sensitivity of the strains indicated to ICP2 phage (left) and quantification of the results from 4 biological replicates (right). **(F)** Phage sensitivity of the strains indicated to the mutant ICP3 phage (left) and quantification of the results from 3-5 biological replicates (right). Data are shown as the standard error of the mean. Analysis was performed using one-way ANOVA with Dunnett’s multiple comparison test (*\*P<0.05, **P<0.01, ***P<0.001, ****P<0.0001*).

### ICP1_2001 encodes proteins that antagonize Nezha

The ICP1 phages used in this study differ across their ∼130 kb genomes by many single nucleotide polymorphisms (SNPs) and a large number of unique regions, making it challenging to identify the genetic difference in ICP1_2001 that counters Nezha. To map the region responsible, we used a phage recombination experimental approach between ICP1_2001 and ICP1_2004_A. ICP1_2004_A was selected because it contains a CRISPR/Cas system with spacers targeting PLE2, thus providing a selection for recombinants of this phage. We reasoned that only recombinant phages that acquire CRISPR/Cas from ICP1_2004_A and the anti-Nezha locus from ICP1_2001 could plaque on a classical A103 derivative modified to contain PLE2. We generated a PLE2 transducing phage to move PLE2 tagged with a Spectinomycin-resistance cassette into wild type A103, Δisland, and *ΔSIR2 ΔherA* derivatives. After isolating transductants, we confirmed that they had the correct phage sensitivity profiles: A103 PLE2 was resistant to both parental phages, while the Δisland PLE2 and *ΔSIR2 ΔherA* PLE2 strains were resistant to ICP1_2001 but sensitive ICP1_2004_A, showing that PLE2 was functional against ICP1_2001 **(Supplemental Figure 3)**.

We generated seven independent recombinant phage pools by co-infecting the permissive El Tor strain E7946 with a 1:1 mixture of ICP1_2001 and ICP1_2004_A at a high multiplicity of infection (MOI) of 5. Recombinant phages that could plaque on the A103 PLE2 host were readily obtained from each pool (example shown in **Figure 4B**). Two phages from each selection were plaque-purified on the A103 PLE2 host and subjected to whole genome sequencing. As expected, mapping the sequencing reads to the ICP1_2004_A parent genome showed the presence of CRISPR/Cas with spacers against PLE2 in all 14 isolates.

**Figure 4.**
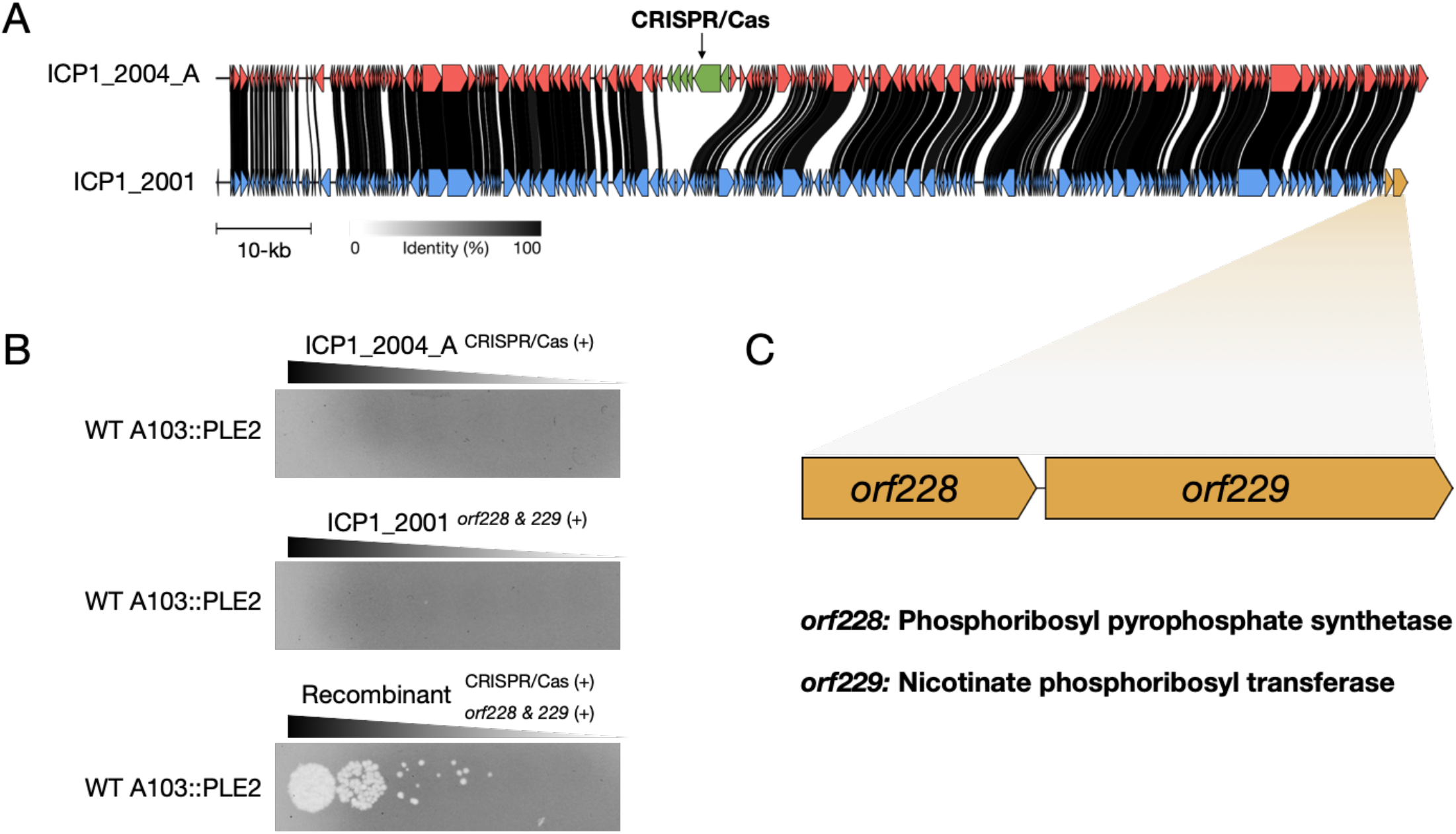
ICP1_2001 encodes for anti-Nezha proteins that overcome NAD^+^ depletion. **(A)** Genome comparison between ICP1_2001 (blue) and ICP1_2004_A (red). Homologous genes are connected with the black lines, where black lines indicate a higher percentage of identity. The CRISPR/Cas system of ICP1_2004_A is shown in green while the anti-Nezha genes of ICP1_2001 are shown in orange. **(B)** Phage sensitivity of classical biotype strain A103::PLE2 to both parent phages and a representative selected recombinant phage. The genotype of the phages is shown. **(C)** Schematic of the anti-Nezha genes encoded by ICP1_2001 and their annotation of the proteins they encode for.

Conversely, mapping the sequencing reads to the ICP1_2001 parent genome showed the presence of the two gene locus located at the 3’ end of the genome in all 14 isolates. The genes in this putative two-gene operon, *orf228* and *orf229,* are annotated as phosphoribosyl pyrophosphate synthetase (PRS) and nicotinate phosphoribosyl transferase (NAPRT), respectively **(Figure 4C)**. It has been shown that PRS uses ribose-5-phosphate and ATP to generate phosphoribosyl pyrophosphate (PRPP). PRPP is utilized by several biosynthetic enzymes, including NAPRT, to generate nicotinic acid mononucleotide, one of several primary substrates that can participate in the synthesis of NAD^+^ (26). Based on Nezha’s anti-phage activity mechanism, we reasoned that ORF228 and ORF229 participate in the regeneration of NAD^+^ within the cell to counter Nezha.

### The ICP1_2001 NAD^+^ regenerating proteins are required to subvert Nezha

After uncovering the potential anti-Nezha genes of ICP1_2001, we wanted to see if other Nezha-sensitive ICP1 isolates lack these two genes. Surprisingly, a Nezha-sensitive isolate, ICP1_2001_A, encodes versions of *orf228* and *orf229*. However, comparing their amino acid sequences reveals that the ICP1_2001_A ORF228 has two amino acid substitutions, V240I and E254G, and ORF229 has an E75G amino acid substitution and a nonsense mutation near the N-terminus (Y29*) that truncates the protein, presumably ablating its function in NAD^+^ regeneration. To test if this defect can be bypassed, we cloned a functional *orf229* from ICP1_2001 into an arabinose inducible expression plasmid, pDL1530, such that ORF229 production is under the control of the P_BAD_ promoter. An empty vector (EV) was included as a control **(Figure 5A).** After mating the plasmids into A103, we induced expression of ORF229 by adding arabinose to the soft agar overlay used for plaque assay. We used wild type A103 and Δisland as controls to show that ICP1_2001_A can only form plaques on the Δisland strain **(Figure 5B)**. Without arabinose, ICP1_2001_A could not form plaques on A103 (pDL1530::*orf229*). However, in the presence of arabinose, we observed complete rescue of ICP1_2001_A plaque formation, with comparable plaquing efficiency on the Δisland strain. As expected, the empty vector control strains were resistant to the phage. This result indicates that an intact *orf229* is needed to counter Nezha and suggests that the variant *orf228* allele in ICP1_2001_A is functional if it plays a role in countering Nezha. To confirm the role of *orf228* in countering Nezha, we did the same experiment using ICP1_2004_A as the infecting phage, which lacks *orf228* and *orf229*. ICP1_2004_A could not form plaques on the A103 strain expressing *orf229* **(Figure 5C)**. This indicates that ORF228 is needed in conjunction with ORF229 to counter Nezha.

**Figure 5.**
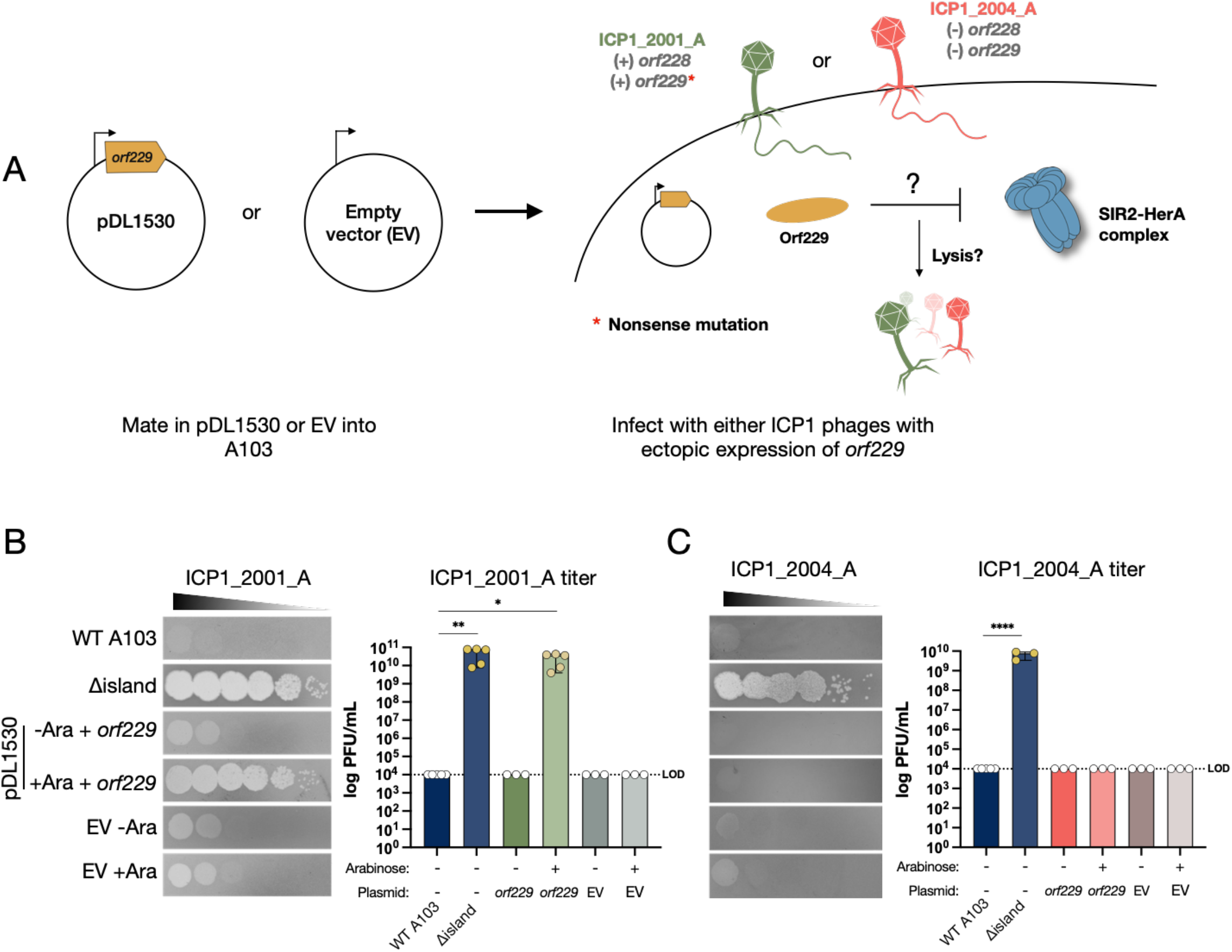
Expressing orf229 in trans rescues ICP1_2001_A but not ICP1_2004_A. **(A)** Cartoon schematic of the experimental outline to test the outcome of orf229 ectopic expression during phage infection with ICP1_2001_A (green) and ICP1_2004_A (red). The genotypes of both phages is shown. **(B)** Phage sensitivity of the strains indicated to ICP1_2001_A or ICP1_2004_A **(C)**. Quantification of the results from 3-5 biological replicates is shown to the right in each panel. Expression of *orf229* from the plasmid was induced by adding arabinose (Ara) to the soft agar overlays. Data are shown as the standard error of the mean for parametric data or as the median with range for non-parametric data. Analysis was performed using one-way ANOVA with Dunnett’s multiple comparison test or Kruskal-Wallis with Dunn’s multiple comparison test (*\*P<0.05, **P<0.01, ***P<0.001, ****P<0.0001*).

Finally, we wanted to test if correcting the nonsense mutation in *orf229* restores the ability of ICP1_2001_A to infect wild-type A103. To do this, we cloned a 338 bp region of the functional *orf229* from ICP1_2001 into pDL1531. After mating the plasmid into the permissive El Tor strain E7946, we passaged ICP1_2001_A twice on this strain. In doing so, we expected some phages to recombine with the plasmid, thus gaining a functional copy of *orf229* **(Figure 6A)**. Recombinant phages were selected on A103, and four plaques were subsequently plaque-purified and subjected to whole genome sequencing. All four had the nonsense mutation corrected to the wild-type tyrosine in addition to reversion of G75E to the wild-type sequence in ICP1_2001. This reversion occurred spontaneously as it was just outside the region cloned into pDL1531. Therefore, this residue may be important for the proper function of ORF229. These results show the importance of encoding a functional ORF228 (PRS) and ORF229 (NAPRT) for ICP1 to counter the Nezha phage defensive system in classical biotype *V. cholerae.* Based on our results and the strong homology of ORF228 and ORF229 to enzymes involved in NAD^+^ synthesis, we named the ICP1 system **<u>N</u>**AD+ **<u>r</u>**egenerating **<u>s</u>**ystem AB (*nrsAB*), where *orf228* is *nrsA* and *orf229* is *nrsB*.

**Figure 6.**
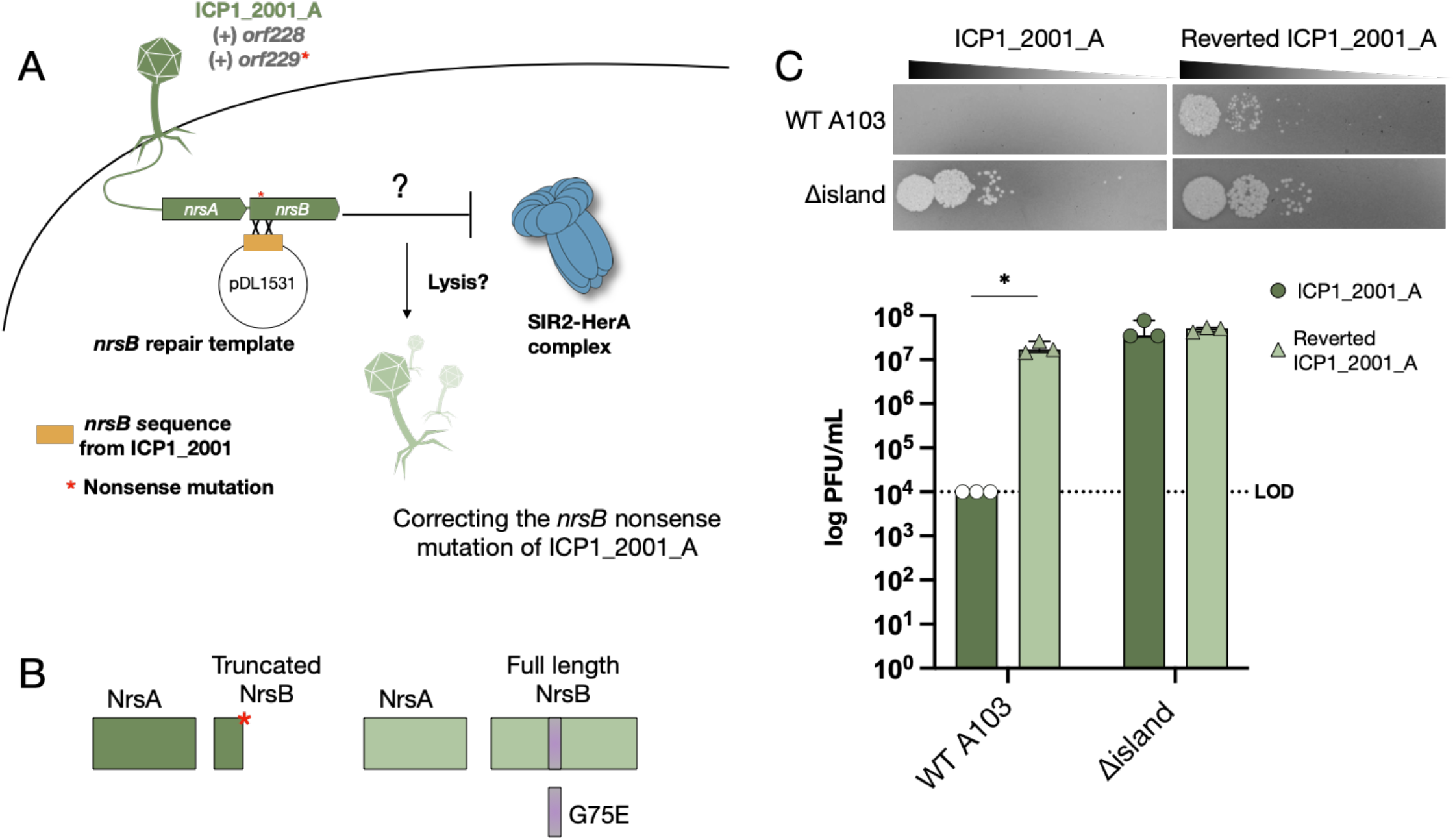
Reverting the nonsense mutation in nrsB rescues ICP1_2001_A infectivity. **(A)** Cartoon schematic of the experiment to revert the nonsense mutation. **(B)** Cartoon schematic of the NrsAB proteins from ICP1_2001_A (left) and NrsAB from reverted ICP1_2001_A (right). **(C)** Phage sensitivity of the strains indicated to ICP1_2001_A or reverted ICP1_2001_A (top) and quantification of the results from 3 biological replicates (bottom). Data are shown as the median with range. Analysis was performed using a Mann-Whitney U test (\**P<0.05*).

## Discussion

In the molecular arms race between bacteria and phages, many anti-phage systems have evolved, which phages evolve to inhibit or circumvent. We identified two anti-phage systems in the classical biotype *V. cholerae*, Gabija and Nezha. Gabija is a high molecular weight complex comprising multiple GajA and GajB subunits (14, 15, 27). GajA is a class 2 OLD (overcoming lysogenization defect) family DNA endonuclease, and GajB belongs to the 1A superfamily of helicase proteins. The role of Gabija in phage defense was first shown using a heterologous system, whereby Gabija from the Gram-positive spore former *Bacillus cereus* was expressed in *E. coli* and shown to provide resistance to T7 phage (15). Gabija systems are present in about 15% of sequenced bacterial and archaeal genomes (13, 20, 28). Similarly, Nezha also encodes for a high molecular weight complex comprising twelve SIR2-like protein (SIR2) subunits and six HerA subunits. SIR2 functions as an NAD^+^ hydrolase with homology to eukaryotic sirtuin proteins (29), while HerA belongs to the helicase family (16, 17). The role of SIR2-like proteins in phage defense was also first shown using a heterologous system, whereby the Thoeris system from *B. cereus* was expressed in *Bacillus subtilis* and shown to provide resistance to several *B. subtilis* phages via NAD^+^ hydrolysis (30, 31). Subsequent studies report that bacteria encode a diverse repertoire of anti-phage defensive systems based on NAD^+^ depletion, including argonautes (pAGO), defense-associated sirtuins (DSR), SEF/IL-17 Receptor (SEFIR), and antiviral ATPase/NTPase of the STAND superfamily (AVAST) (19, 32, 33)

In contrast to many of the studies cited above that used heterologous systems to show the function of Gabija and Nezha in phage defense, we show their function in a natural host/phage system. Using a series of defined deletions, we showed that in classical biotype *V. cholerae*, Gabija plays a minor role in defense against ICP1 and ICP2 phages and that Nezha plays a major role in resistance to ICP1, ICP2, and ICP3. We confirmed this by mutating the active site of SIR2 to abolish NAD^+^ hydrolase activity, which resulted in a complete loss of phage resistance. Moreover, deleting the HerA helicase resulted in the loss of phage resistance. This is consistent with a previous report that the helicase subunit of Nezha is required for the NADase activity of SIR2 (16, 17).

Nezha hydrolyzes NAD^+^ to generate adenosine diphosphate ribose (ADPR) and nicotinamide (NAM), thus depleting this essential coenzyme within the host cell. In a *B. subtilis* strain engineered to express a homologous DSR2 system from another *B. subtilis* strain, NAD^+^ depletion by DSR2 blocked SPR phage replication, while phi3T phage was resistant to the effects of DSR2. phi3T encodes for an anti-DSR2 protein called DSAD1, which directly interacts with and antagonizes the NAD^+^-depleting effector (19). In contrast, we showed that phage ICP1_2001 indirectly counters the effect of Nezha by encoding two putative NAD^+^ synthesis pathway genes, *nrsA* and *nrsB*.

Of the 75 ICP1 isolate sequences deposited in Genbank, five lack *nrsAB,* and 60 have the Y29* mutation in *nrsB*. Functional *nrsAB* sequences are only present in four strains isolated from Bangladesh before 2002 and six strains isolated from the Congo in 2017. If *V. cholerae* in those locations at those times contained Nezha (or another NADase) there would have been a selection to maintain functional *nrsAB* genes. In contrast, although Nezha is found in some current environmental strains of *V. cholerae*, it is not found in hundreds of pathogenic strains isolated from the stools of cholera patients. Hence, for most ICP1 phages, there may be no selection to maintain functional *nrsAB* genes. Moreover, if this operon imposes a fitness cost to ICP1 by interfering with host NAD^+^ metabolism during phage infection, it would be advantageous to lose or inactivate the *nrsAB* genes. Consistent with this, while we could clone and express *nrsB* alone in *V. cholerae*, we could not do so with *nrsAB,* suggesting it is toxic.

Based on homology, we initially assumed NrsA was a phosphoribosyl pyrophosphate synthetase, converting ribose 5-phosphate to phosphoribosyl pyrophosphate, and NrsB, a nicotinate phosphoribosyl transferase that adds nicotinic acid to phosphoribosyl pyrophosphate to generate nicotinic acid mononucleotide (34). However, during the preparation of this manuscript, a biochemical characterization of proteins highly homologous to NrsAB was deposited in bioRxiv (35). The authors reported that homologs of NrsA and NrsB are adenosine diphosphate ribose-pyrophosphate synthetases (Adps) and nicotinamide ADPR-transferases (Namat), respectively. It was shown that Adps adds a pyrophosphate (PP) to ADPR in an ATP-dependent manner to generate ADPR-PP. This molecule is then utilized by the second enzyme, Namat, to catalyze a previously uncharacterized reaction that conjugates nicotinamide onto ADPR-PP, directly generating NAD^+^. Given that ICP1 NrsA and NrsB share strong structural homology over their entire lengths with Adps and Namat, including identity across the active site residues **(Supplemental Figures 4 and 5)**, NrsA and NrsB likely catalyze the same reactions, thus comprising an NAD^+^ regenerating system.

In summary, we establish that the classical biotype *V. cholerae* encodes two phage defensive systems, Gabija and Nezha, absent from El Tor biotype strains. Nezha is the more robust defense against the ICP phages. Some ICP1 phages harbor a functional NAD^+^ regenerating system to counter Nezha and other NADases, while the majority harbor a non-functional system that can be easily reactivated by a reversion mutation should NADases be encountered. This adds to the rapidly advancing knowledge of the diverse molecular mechanisms bacteria and phages use against each other. This is important not only for understanding the impact of phages on the life cycles and evolution of bacteria but also informs the use of phages in preventing and treating bacterial infections.

## Materials and Methods

### Bacterial strains and growth conditions

Bacterial strains, phages, and plasmids used in this study are listed in Table 2. The complete, annotated plasmid sequences in Genbank format are provided in **Supplementary Files 1, 2, and 3**. Bacteria were cultured at 37°C in Lysogeny broth (LB) with aeration or on LB agar. Media were supplemented with 50 µg/mL of Apramycin (Apra), Spectinomycin (Spec), Kanamycin (Kan), Carbenicillin (Carb), or 100 µg/mL Streptomycin (Sm) when appropriate. A diaminopimelic acid (DAP) auxotrophic Pir- expressing RP4 conjugation donor derivative of *E. coli* TG1, called TG1 donor pir, was constructed in two steps. First, we used P1 to transduce the RP4 mating function genes flanked by Apramycin- and Zeocin-resistance genes from *E. coli* MFDpir (36) into TG1 to create TG1 donor. TG1 donor pir was then created using lambda-red recombineering with a PCR fragment amplified from MFDpir in which the dapA gene was replaced with those that express pir and erythromycin resistance. TG1 donor pir strains were supplemented with 0.5 mM DAP. Plasmids pDL1530 and pDL1531 were constructed for this study to allow inducible expression of cloned genes and for allelic exchange, respectively. Both have an p15a origin of replication and an RP4 oriT for conjugal transfer. Their complete annotated sequences are provided in Supplemental file 2. Plasmids were induced with either 1 mM isopropyl-ß-D- thiogalactopyranoside (IPTG) or 0.2% arabinose (Ara).

### Phage propagation

ICP1, ICP2, and ICP3 phages (Table 2) were those originally isolated from cholera patient rice-water stool samples in Dhaka, Bangladesh (insert citation). All phage mutants constructed in this study were plaque-purified three times. High titer stocks were made by infecting mid-exponential growth phase cultures of *V. cholerae* E7946 or derivatives grown in LB at 37°C at an MOI of 0.01 and then incubating the cultures for 2-4 hrs at 37°C with aeration. Infected cultures were chilled to 4°C, centrifuged at 10,000 RCF for 15 mins at 4°C, and the supernatants were filter-sterilized through 0.45 µm bottle-top filters. Phages were concentrated by precipitation overnight at 4°C after adding 1x polyethylene glycol (4% PEG 8000, 0.5M NaCl). The precipitated phages were centrifuged at 15,000 RCF for 10 mins at 4°C, and the supernatant removed. The phage pellets were resuspended in phage 80 buffer (0.1mM MgSO4, 0.1M Tris-HCl pH 7.4, 85.6mM NaCl) or 0.7% (w/v) Instant Ocean salts (IO).

### Bacterial conjugation

Chemically competent *E. coli* TG1 donor pir cells were first transformed with the plasmids listed in Table 2. Next, plasmids were moved into *V. cholerae* by mating with TG1 donor pir cells. Briefly, donor and recipient cells were cultured overnight in LB supplemented with DAP and Carb or LB alone at 37°C, respectively. The next day, 0.5 ml of donor and recipient were pelleted and washed twice with LB before resuspending in 50 µL of LB. A 1:1 mixture was added to a sterile 0.2 µm filter (Millipore) on an LB plate supplemented with DAP and incubated at 37°C for 3 hrs. Cells were recovered from the filter by vortexing in 1 ml LB. Serial dilutions were plated on LB agar supplemented with Carb to select for the exconjugates.

#### Construction of *V. cholerae* mutants

Marked deletion and point mutation constructs were made by splicing-by-overlap extension (SOE) PCR, and then the linear dsDNA products were transformed into naturally competent *V. cholerae*. Natural competence was induced by growth overnight in IO containing shrimp chitin flakes at 30°C without aeration. Transformants were selected on LB agar supplement with the appropriate antibiotic(s).

### Plaque assays

Bacterial strains were grown overnight in LB at 37°C with aeration. The cultures were then back-diluted into fresh LB and grown at 37°C to an OD_600_ of 0.5. 0.1 mL of the culture was added to 6 mL of 50°C soft agarose (LB with 0.3% agarose) and poured on top of LB agar plates and allowed to cool and solidify. Phage were serially diluted ten-fold and 5 µL was spotted or 10 µL was dribbled on the overlay. Plates were incubated at 37°C for 3-4 hrs, and the number of plaques counted. For NrsB complementation, A103 (pDL1530) was grown in LB supplemented with Kan to an OD_600_ of 0.5. 0.1 mL of the culture was added to 6 mL of 50°C soft agarose supplemented with Kan and Ara to maintain the plasmid and induce expression of the P_BAD_ promoter, respectively.

### Transduction of PLE2

Natural competence was used to insert the aad9 gene flanked by promoter and terminators downstream of and in the same direction as the orf2 gene in PLE2. The resulting strain was then infected with ICP1_2001 at a MOI of 1. After 2 hrs of incubation at 37°C, the infected culture was centrifuged, and the supernatant was filtered. The recipient A103 was grown overnight at 37°C in LB. The next day, 0.1 mL of the overnight culture was infected with 10 µL of the filtrate and the volume was adjusted to 1 mL by addition of 0.89 mL of LB. The culture was incubated for 1 hr at 37°C and plated on LB agar supplemented with Spec to select for transductants.

### Generating recombinant phage

E7946 was grown overnight in LB at 37°C and back diluted the next day by adding 0.1 mL into seven culture tubes each containing 10 mL LB. The culture tubes were incubated at 37°C for 20 mins with aeration and then co-infected with a 1:1 mixture of ICP1_2001 and ICP1_2004_A at an MOI of 5. After incubating at 37°C for 2 hrs, the infected culture was centrifuged at 10,000 RCF for 15 mins and the supernatant was filtered. Phage were plaqued on A103 PLE2::*aad9* to isolate recombinants and passaged three times. Agar stabs with phage were isolated to generate high titer recombinant phage stocks using E7946. Phage DNA was isolated from a portion of the high titer stocks for whole genome sequencing.

### Whole genome sequencing

Genomic DNA was extracted from bacteria or phage using the Zymo gDNA prep kit (Zymo Research, Corp) or the Promega Wizard kit (Promega, Inc), respectively, following the manufacturer’s protocol. Most bacterial and phage samples were sequenced using single or paired-end sequencing on the Illumina Nextseq platform (Illumina, Inc). ICP3 mutants were sequenced using Plasmidsaurus (Plasmidsauraus, Inc).

### Statistical analysis

Data were first log-transformed and checked for normality using the Shapiro-Wilk test. Parametric data were analyzed using the one-way ANOVA test with Dunnett’s multiple comparisons. Non-parametric data were analyzed using either the Mann-Whitney U test or the Kruskal-Wallis test with Dunn’s multiple comparisons. P values <0.05 were considered as significant. GraphPad Prism software, version 10, was used for all statistical analysis.

## SUPPLEMENTARY MATERIALS

**Supplementary Figure 1.**
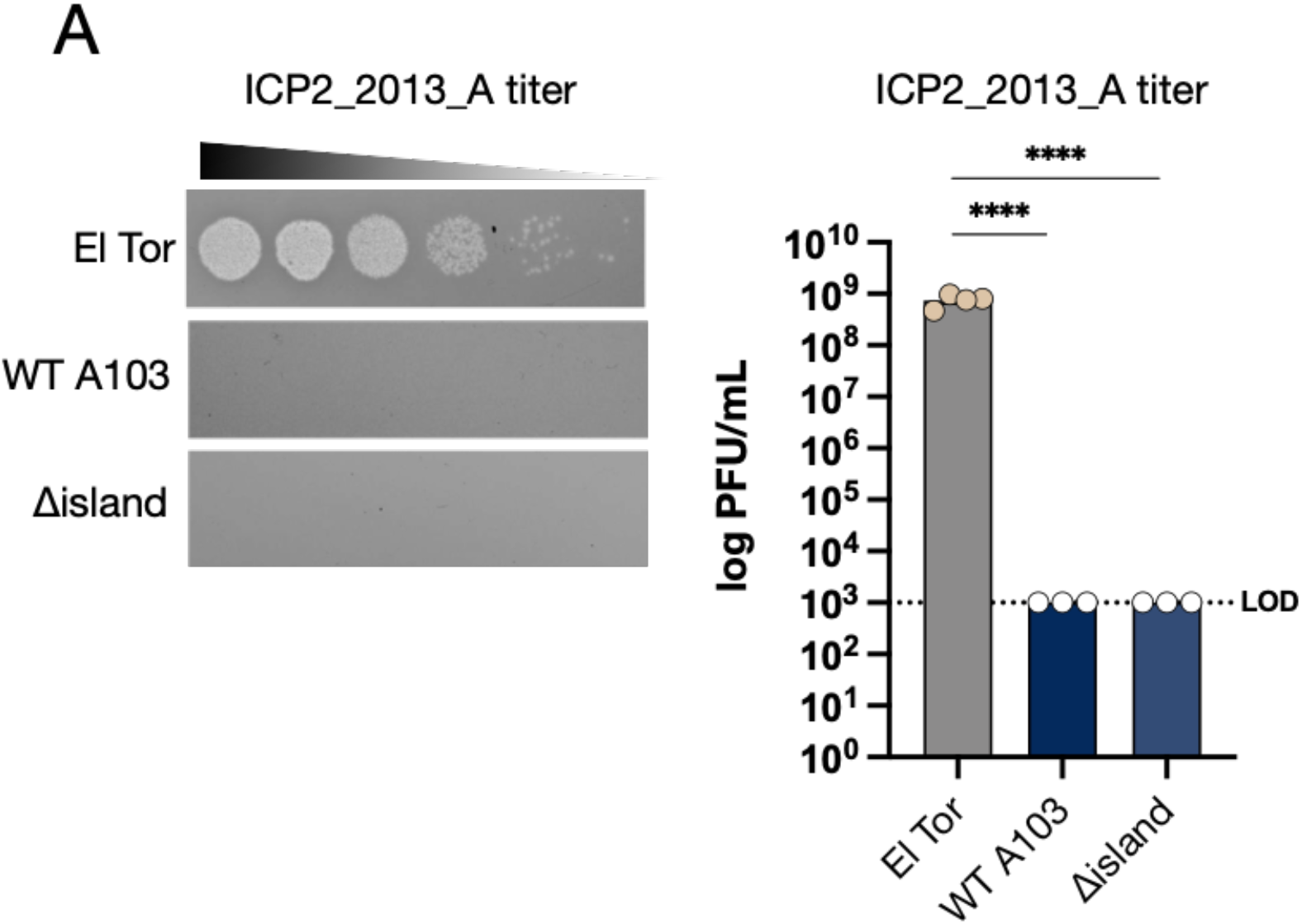
A103 and its derivative Δisland are resistant to ICP2 due to an OmpU mutation. **(A)** Phage sensitivity of the strains indicated to phage ICP2_2013_A (left) and quantification of the results from 3-4 biological replicates (right). El Tor is strain E7946; the rest are classical strain A103 wild type and derivatives. The two classical strains have a mutated OmpU receptor which prevents WT ICP2 infection. Data are shown as the standard error of the mean. Analysis was performed using one-way ANOVA with Dunnett’s multiple comparison test (*\*P<0.05, **P<0.01, ***P<0.001*, ****P<0.0001).

**Supplementary Figure 2.**
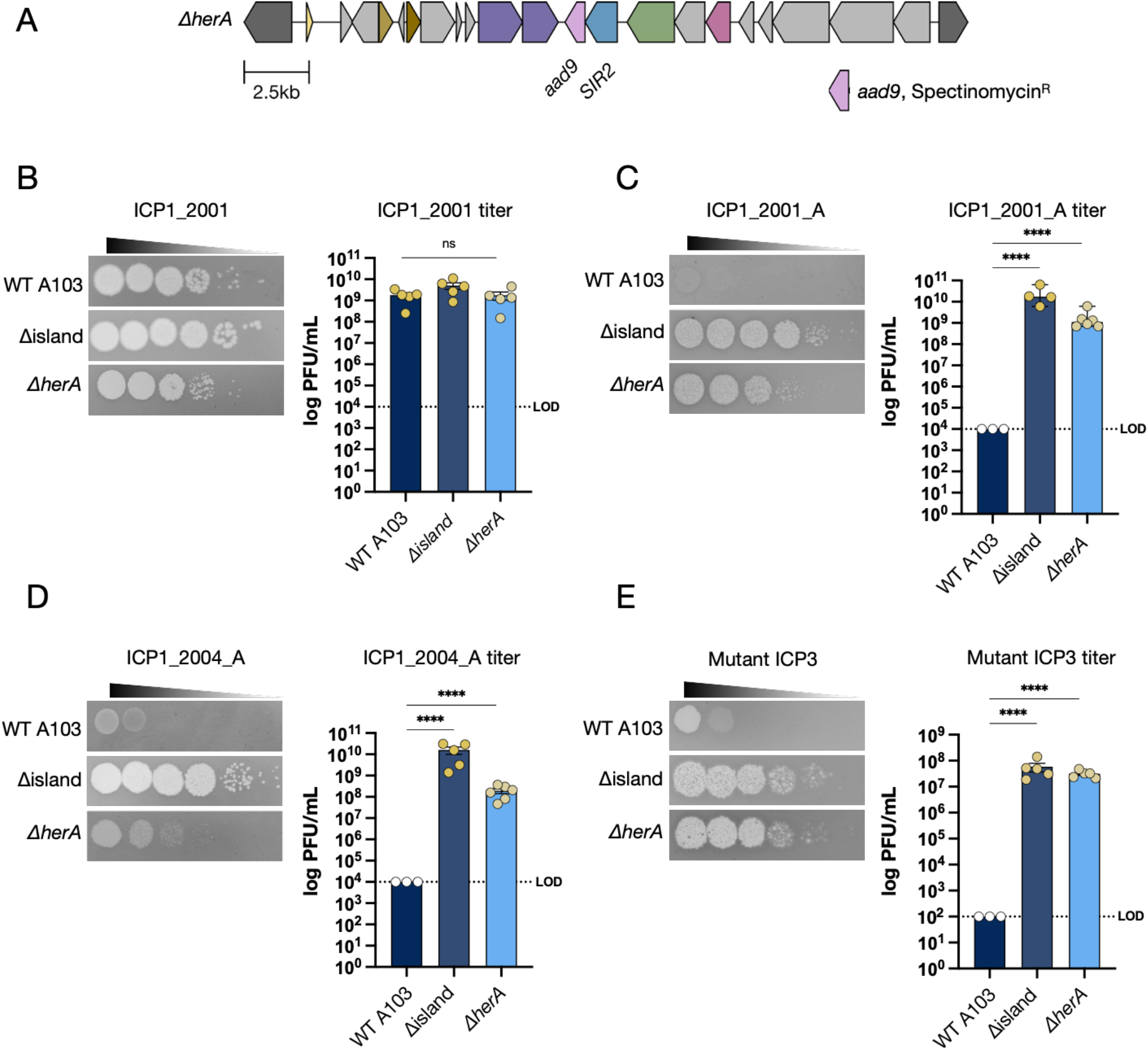
Deletion of *herA* abolishes immunity to ICP1, ICP2, and ICP3. **(A)** Schematic of the *ΔherA* deletion construct, where *herA* was replaced with a spectinomycin-resistance cassette (pink). (**B-D)** Phage sensitivity of the strains indicated to ICP1 phages (left) and quantification of the results from 3-5 biological replicates (right). **(E)** Phage sensitivity of the strains indicated to ICP3 phage (left) and quantification of the results from 3-5 biological replicates (right). Data are shown as the standard error of the mean for normally distributed data or as the median with range for non-normally distributed data. Data are shown as the standard error of the mean. Data were analyzed using one-way ANOVA with Dunnett’s multiple comparison test (*\*P<0.05, **P<0.01, ***P<0.001, ****P<0.0001*).

**Supplementary Figure 3.**
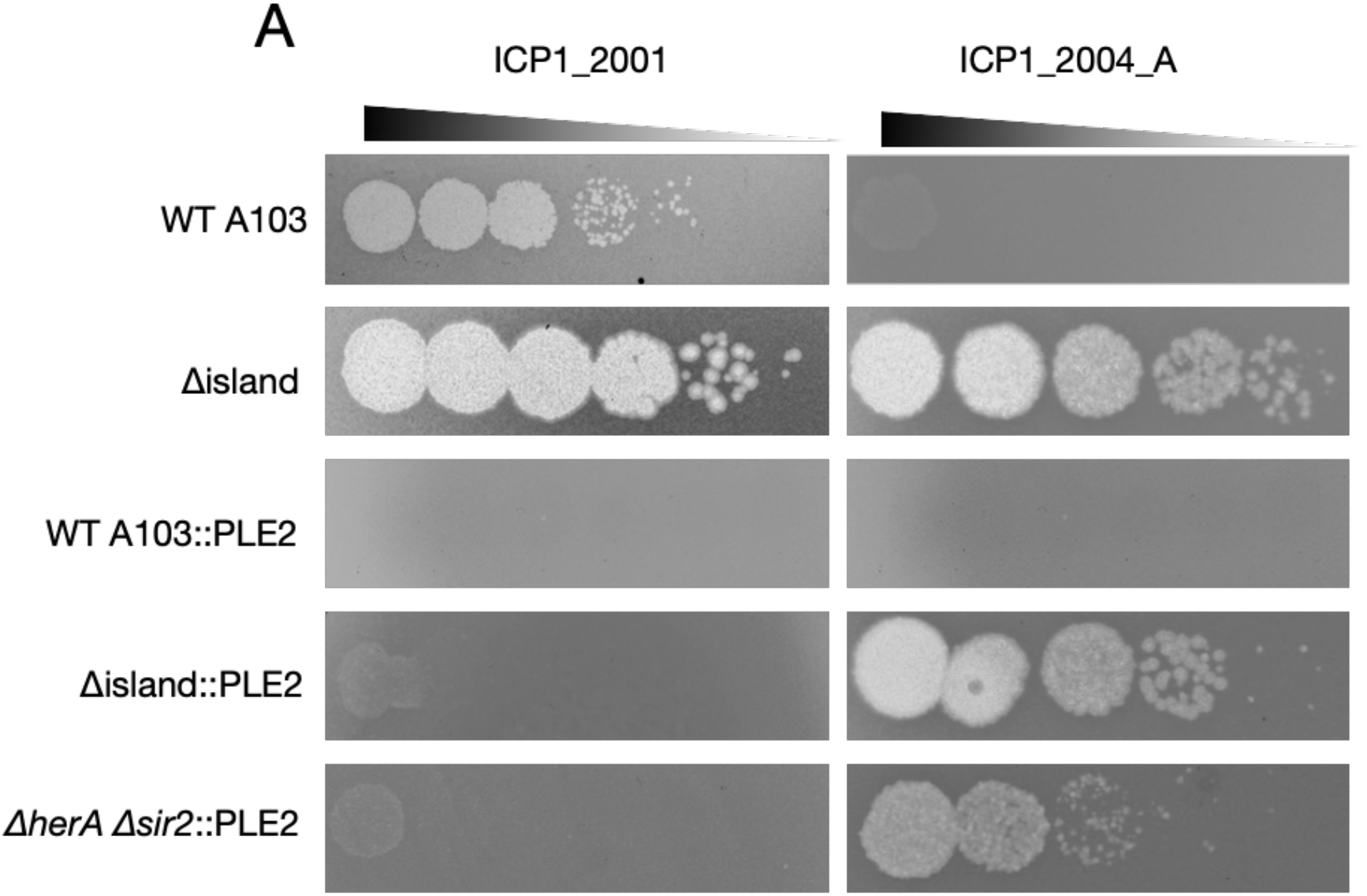
PLE2 antagonizes ICP1_2001 but not ICP1_2004_A. **(A)** Phage sensitivity of classical biotype strain A103 and its derivatives containing PLE2 against ICP1_2001 or ICP1_2004_A. PLE2 is a potent inhibitor of ICP1_2001, but not ICP1_2004_A, which contains a CRISPR/Cas system that targets PLE2.

**Supplementary Figure 4.**
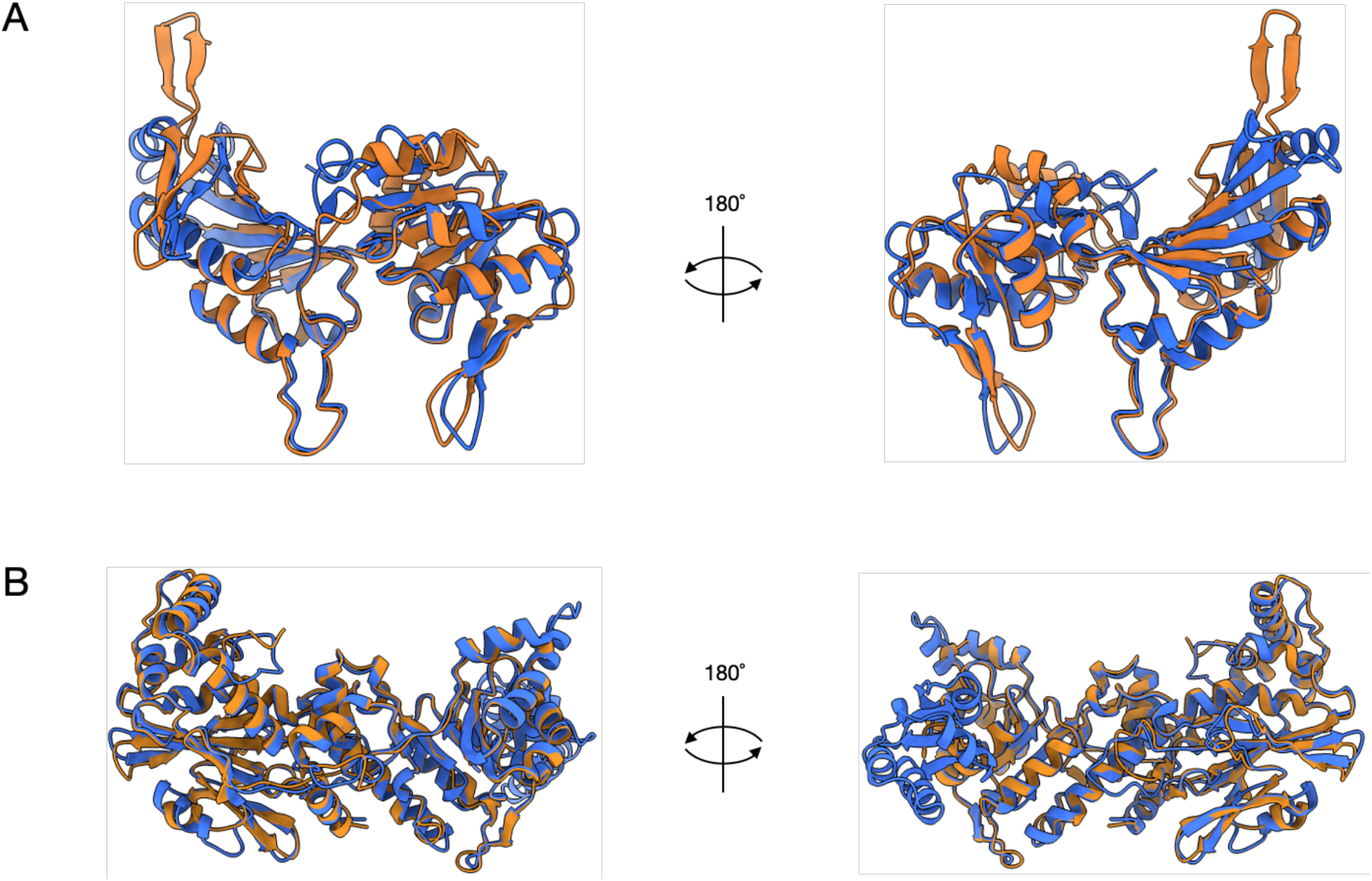
Structural alignment of ICP1 and Bas63 NAD^+^ regenerating proteins. **(A)** Alpha fold protein prediction of ICP1_2001 NrsA (orange) overlayed with Bas63 Adps (blue). The structures were rotated 180° to visualize the proteins from the back. **(B)** Alpha fold protein prediction of ICP1_2001 NrsB (orange) overlayed with Bas63 Namat (blue). The structures were rotated 180° to visualize the proteins from the back.

**Supplementary Figure 5.**
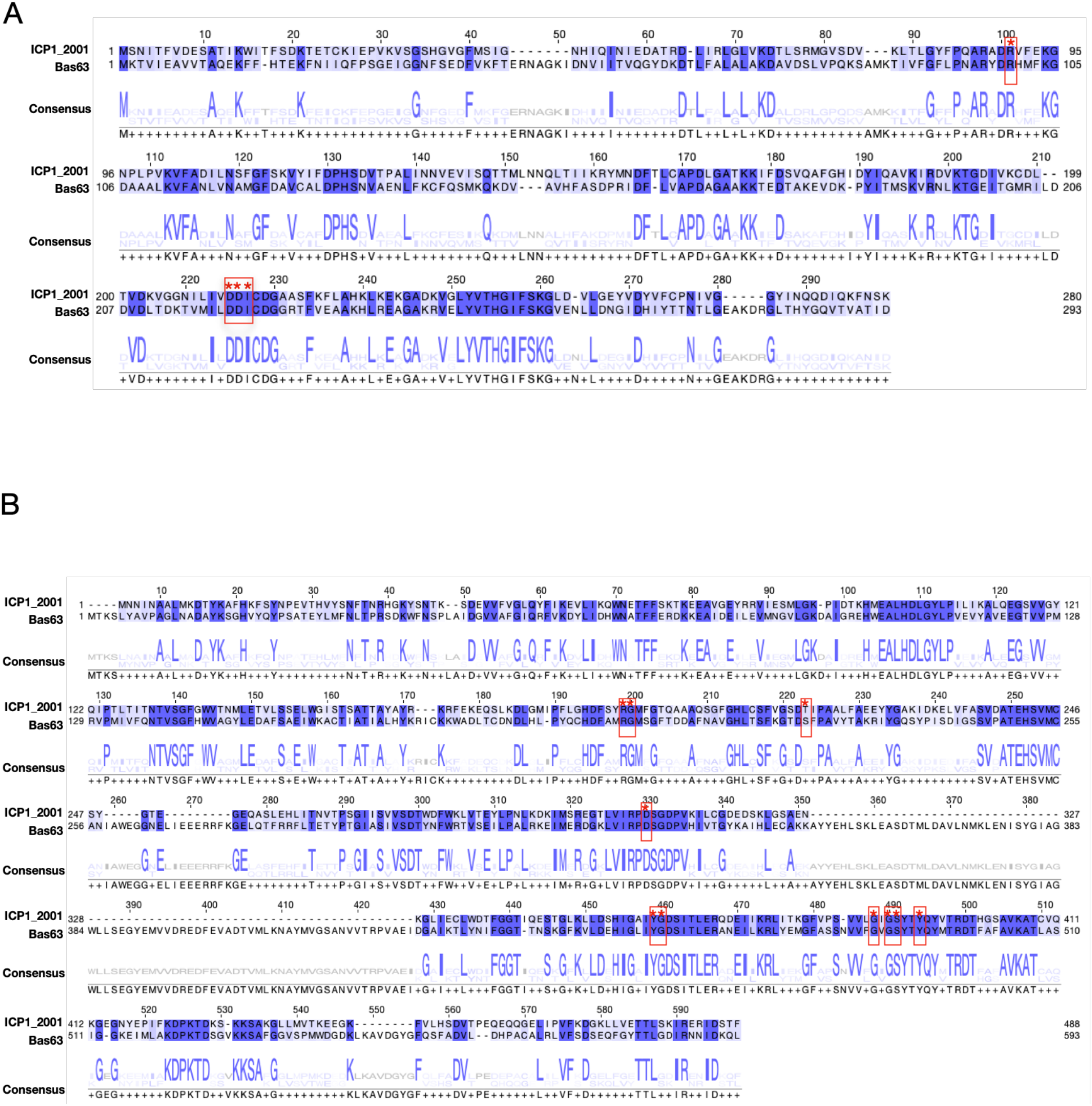
Comparing the ICP1_2001 ORF228 and ORF229 with homologues from *E. coli* phage Bas63. **A)** Protein alignment using the MUSCLE algorithm of ICP1_2001 ORF228 (top) and Bas63 0079 (bottom). Red boxes indicate active site residues that are conserved between the two proteins. **B)** Protein alignment using the MUSCLE algorithm of ICP1_2001 ORF229 (top) and Bas63 0080 (bottom). Red boxes indicate active site residues that are conserved between the two proteins.

**Supplemental Table 1.** The ICP3 mutant phage has a frame-shift mutation in *orf17*.

| Accession # | Gene annotation | Coding region change | Amino acid change |
| --- | --- | --- | --- |
| **NC_015159** | Mutant 1:*orf17* | YP_004251260.1: c.74del T | YP_004251260.1: p.Phe26 fs(frame shift) |
|  | Mutant 2:*orf17* | YP_004251260.1: c.74del T | YP_004251260.1: p.Phe26 fs(frame shift) |
|  | Mutant 3:*orf17* | YP_004251260.1: c.130del A | YP_004251260.1: p.Thr45 fs(frame shift) |
|  | Mutant 4:*orf17* | YP_004251260.1: c.130del A | YP_004251260.1: p.Thr45 fs(frame shift) |

**Supplementary Table 2.**
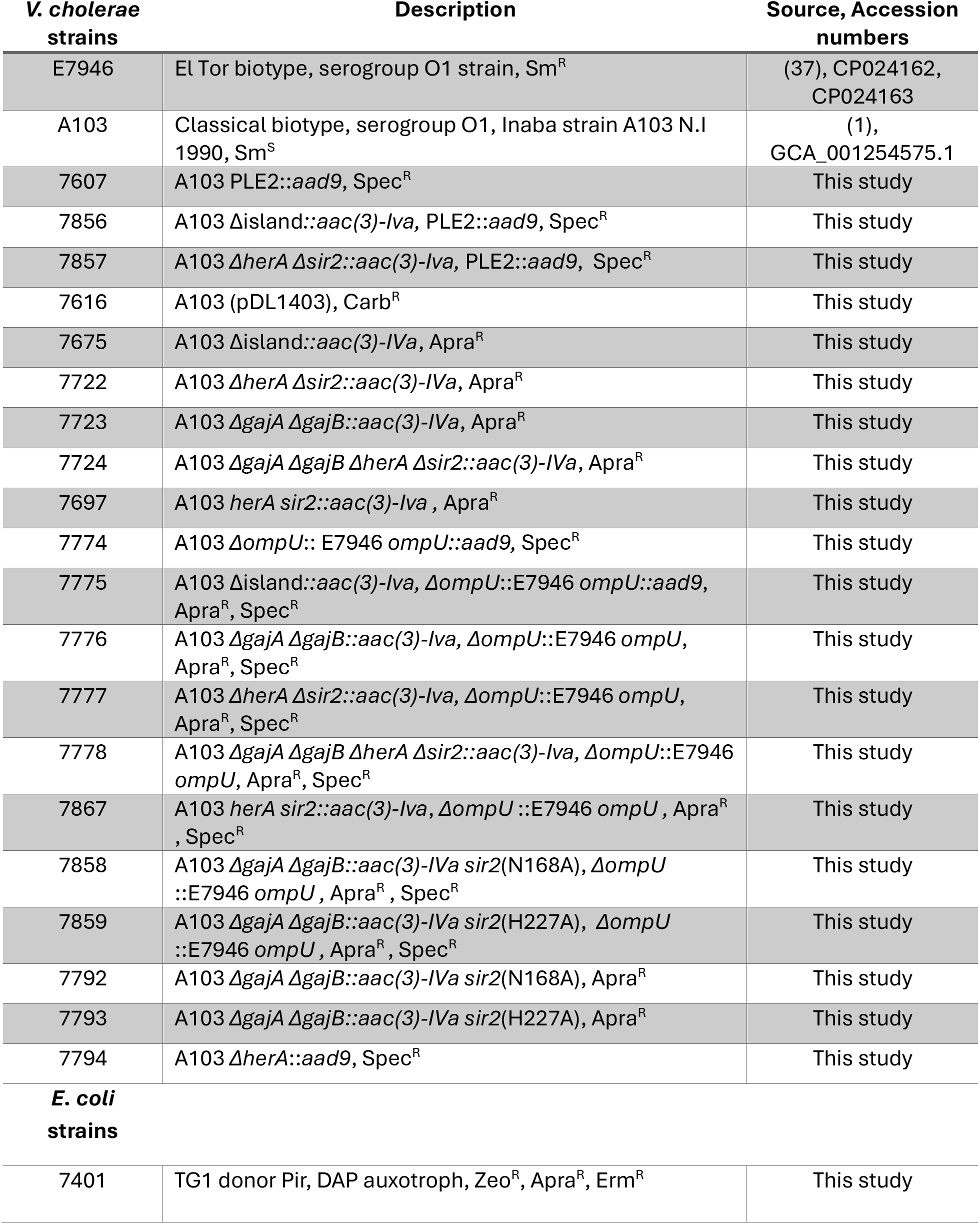
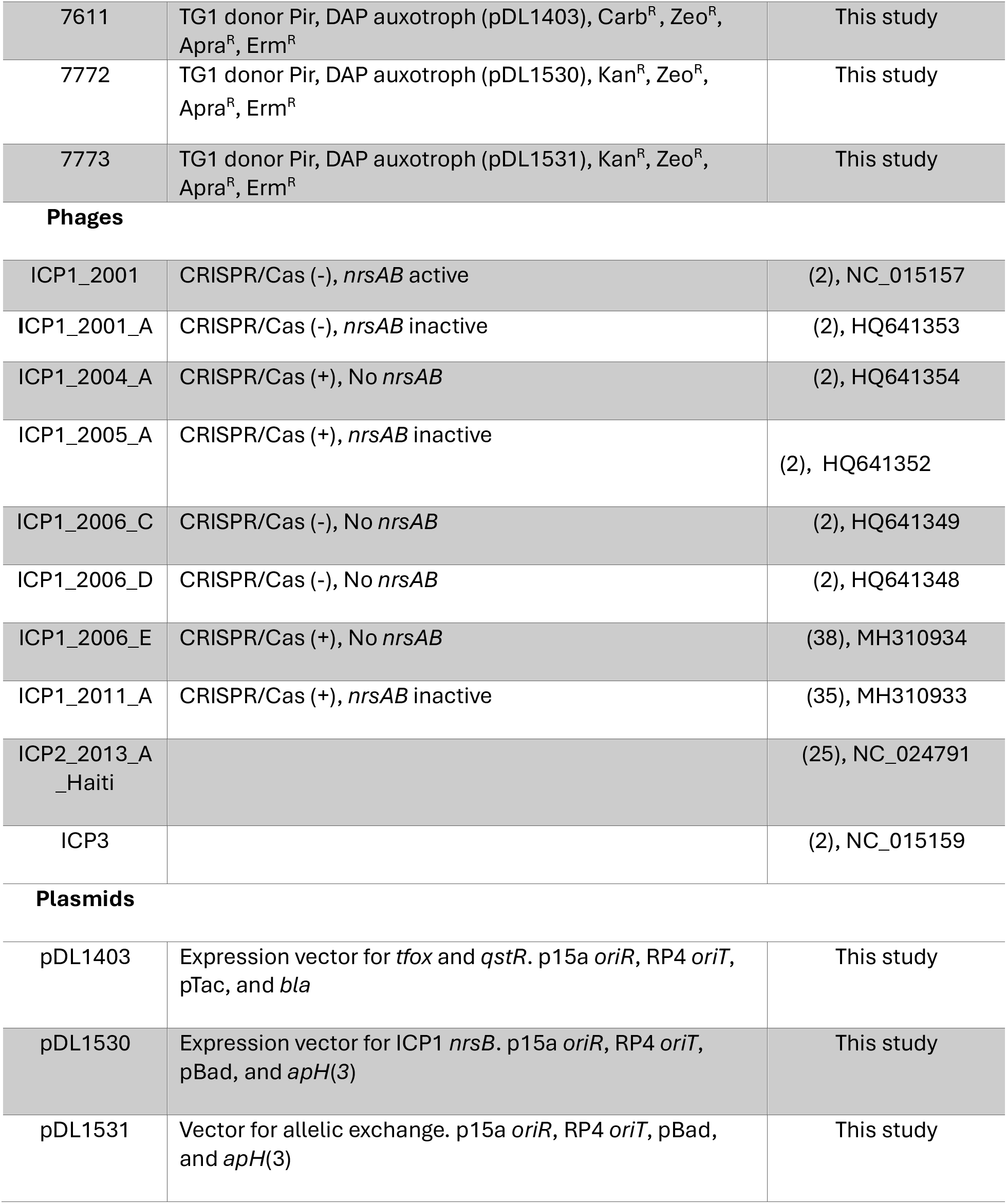
Bacterial strains, phages, and plasmids used.

| ***V. cholerae***<br>**strains** | **Description** | **Source, Accession**<br>**numbers** |
| --- | --- | --- |
| E7946 | El Tor biotype, serogroup O1 strain, Sm<sup>R</sup> | (37), CP024162,<br>CP024163 |
| A103 | Classical biotype, serogroup O1, Inaba strain A103 N.I<br>1990, Sm<sup>S</sup> | (1),<br>GCA_001254575.1 |
| 7607 | A103 PLE2::*aad9*, Spec<sup>R</sup> | This study |
| 7856 | A103 Δisland::*aac(3)-Iva*, PLE2::*aad9*, Spec<sup>R</sup> | This study |
| 7857 | A103 ΔherA Δsir2::*aac(3)-Iva*, PLE2::*aad9*, Spec<sup>R</sup> | This study |
| 7616 | A103 (pDL1403), Carb<sup>R</sup> | This study |
| 7675 | A103 Δisland::*aac(3)-Iva*, Apra<sup>R</sup> | This study |
| 7722 | A103 ΔherA Δsir2::*aac(3)-Iva*, Apra<sup>R</sup> | This study |
| 7723 | A103 ΔgajA ΔgajB::*aac(3)-Iva*, Apra<sup>R</sup> | This study |
| 7724 | A103 ΔgajA ΔgajB ΔherA Δsir2::*aac(3)-Iva*, Apra<sup>R</sup> | This study |
| 7697 | A103 *herA sir2*::*aac(3)-Iva*, Apra<sup>R</sup> | This study |
| 7774 | A103 ΔompU:: E7946 *ompU*::*aad9*, Spec<sup>R</sup> | This study |
| 7775 | A103 Δisland::*aac(3)-Iva*, ΔompU::E7946 *ompU*::*aad9*,<br>Apra<sup>R</sup>, Spec<sup>R</sup> | This study |
| 7776 | A103 ΔgajA ΔgajB::*aac(3)-Iva*, ΔompU::E7946 *ompU*,<br>Apra<sup>R</sup>, Spec<sup>R</sup> | This study |
| 7777 | A103 ΔherA Δsir2::*aac(3)-Iva*, ΔompU::E7946 *ompU*,<br>Apra<sup>R</sup>, Spec<sup>R</sup> | This study |
| 7778 | A103 ΔgajA ΔgajB ΔherA Δsir2::*aac(3)-Iva*, ΔompU::E7946<br>*ompU*, Apra<sup>R</sup>, Spec<sup>R</sup> | This study |
| 7867 | A103 *herA sir2*::*aac(3)-Iva*, ΔompU::E7946 *ompU*, Apra<sup>R</sup><br>, Spec<sup>R</sup> | This study |
| 7858 | A103 ΔgajA ΔgajB::*aac(3)-Iva sir2*(N168A), ΔompU<br>::E7946 *ompU*, Apra<sup>R</sup>, Spec<sup>R</sup> | This study |
| 7859 | A103 ΔgajA ΔgajB::*aac(3)-Iva sir2*(H227A), ΔompU<br>::E7946 *ompU*, Apra<sup>R</sup>, Spec<sup>R</sup> | This study |
| 7792 | A103 ΔgajA ΔgajB::*aac(3)-Iva sir2*(N168A), Apra<sup>R</sup> | This study |
| 7793 | A103 ΔgajA ΔgajB::*aac(3)-Iva sir2*(H227A), Apra<sup>R</sup> | This study |
| 7794 | A103 ΔherA::*aad9*, Spec<sup>R</sup> | This study |
| ***E. coli***<br>**strains** |  |  |
| 7401 | TG1 donor Pir, DAP auxotroph, Zeo<sup>R</sup>, Apra<sup>R</sup>, Erm<sup>R</sup> | This study |
| 7611 | TG1 donor Pir, DAP auxotroph (pDL1403), Carb<sup>R</sup>, Zeo<sup>R</sup>, Apra<sup>R</sup>, Erm<sup>R</sup> | This study |
| 7772 | TG1 donor Pir, DAP auxotroph (pDL1530), Kan<sup>R</sup>, Zeo<sup>R</sup>, Apra<sup>R</sup>, Erm<sup>R</sup> | This study |
| 7773 | TG1 donor Pir, DAP auxotroph (pDL1531), Kan<sup>R</sup>, Zeo<sup>R</sup>, Apra<sup>R</sup>, Erm<sup>R</sup> | This study |
| Phages |  |  |
| ICP1_2001 | CRISPR/Cas (-), *nrsAB* active | (2), NC_015157 |
| ICP1_2001_A | CRISPR/Cas (-), *nrsAB* inactive | (2), HQ641353 |
| ICP1_2004_A | CRISPR/Cas (+), No *nrsAB* | (2), HQ641354 |
| ICP1_2005_A | CRISPR/Cas (+), *nrsAB* inactive | (2), HQ641352 |
| ICP1_2006_C | CRISPR/Cas (-), No *nrsAB* | (2), HQ641349 |
| ICP1_2006_D | CRISPR/Cas (-), No *nrsAB* | (2), HQ641348 |
| ICP1_2006_E | CRISPR/Cas (+), No *nrsAB* | (38), MH310934 |
| ICP1_2011_A | CRISPR/Cas (+), *nrsAB* inactive | (35), MH310933 |
| ICP2_2013_A_Haiti |  | (25), NC_024791 |
| ICP3 |  | (2), NC_015159 |
| Plasmids |  |  |
| pDL1403 | Expression vector for *tfox* and *qstR*. p15a *oriR*, RP4 *oriT*, pTac, and *bla* | This study |
| pDL1530 | Expression vector for ICP1 *nrsB*. p15a *oriR*, RP4 *oriT*, pBad, and *apH(3)* | This study |
| pDL1531 | Vector for allelic exchange. p15a *oriR*, RP4 *oriT*, pBad, and *apH(3)* | This study |

**Supplementary Files 1, 2 and 3**. Annotated plasmid sequences of pDL1403, pDL1530, and pDL1531 in Genbank format.

